# Crystal structures of human SP-D complexed with synthetic oligosaccharides suggest a role for phosphorylated inner core LPS saccharides in host-pathogen interactions

**DOI:** 10.1101/2025.05.08.652840

**Authors:** Harry M. Williams, Alastair Watson, Jens Madsen, Howard W. Clark, Derek W. Hood, Stefan Oscarson, Trevor J. Greenhough, Annette K. Shrive

**Author notes:** To whom correspondence should be addressed: Annette K Shrive, School of Life Sciences, Keele University, Staffordshire, ST5 5BG, UK. Laboratory of Molecular Nanodynamics, Ecole Polytechnique Fédérale de Lausanne (EPFL), CH-1015 Lausanne, Switzerland.

## Abstract

The innate immune protein human surfactant protein D (SP-D) recognises pathogens in the lungs via binding to carbohydrate surface structures. SP-D has been shown to target gram-negative bacterial lipopolysaccharide via calcium-dependent binding, preferentially to the inner core heptose (HepI). To further investigate this recognition, we have determined the high-resolution crystal structures of a trimeric recombinant fragment of human SP-D complexed with synthetic di- and trisaccharides, HepI-Kdo, HepIII-HepII-HepI, and HepII-HepI phosphorylated at either HepI or HepII, inner core motifs common to the lipopolysaccharide of many gram-negative bacteria. In contrast to acid-hydrolysed lipopolysaccharide used in several previous studies, these synthetic saccharides allow presentation of both the innermost Kdo in its natural pyranose form and heptose phosphorylation. The structures confirm the flexibility of SP-D to adopt an alternative binding mode when the preferred epitope is not available, reveal a preference for recognition of the reducing terminal heptose (HepI) via the glyceryl group, indicate that a single Kdo attached to HepI does not have a significant role in ligand recognition, and provide evidence that recognition of phosphorylated inner core diheptosyl ligands varies dependent on whether HepI or HepII is phosphorylated. The disaccharide with HepII O4’ phosphorylation binds via the preferred HepI glyceryl-hydroxyls, while HepI O4’ phosphorylation reveals HepII binding via the pyranose ring O3’ and O4’ hydroxyls. The ability of the HepII O4’ phosphate to prevent preferred HepI recognition suggests a role for heptose phosphorylation in shielding the bacterial LPS inner core from immune recognition.

## INTRODUCTION

Surfactant protein D (SP-D) is a collectin with a key role in the innate immune defence that is expressed in pulmonary, as well as non-pulmonary epithelia (Madsen *et al*., 2000; Sorensen *et al*., 2006; Brauer *et al*., 2009; Schob *et al*., 2013; Rokade; *et al*., 2016). Human SP-D (hSP-D) is known to exert significant antimicrobial effects principally via opsonisation (Restrepo *et al*., 1999; LeVine *et al*., 2004; Tecle *et al*., 2008; Thawer *et al*., 2016) whilst also having a role in modulating the broader host response to pathogens by interacting with various cell-based receptors (Watson *et al*., 2021) including the osteoclast-associated receptor, also known as OSCAR (Barrow *et al*., 2015), DC-SIGN (Dodagatta-Marri *et al*., 2017) and natural killer cell membrane receptor, NKp46 (Ge *et al*., 2016).

The recognition of bacterial, fungal, and viral pathogens by hSP-D is mediated by the carbohydrate recognition domain (CRD), which detects carbohydrate residues embedded within the surface structures of pathogens. For gram-negative bacteria this is achieved through calcium-dependent binding of bacterial lipopolysaccharide (LPS) carbohydrate via a conserved hydroxyl motif. Bacterial LPS is typically composed of three major domains: the lipid A domain, core oligosaccharide, and hypervariable O-polysaccharide (Raetz and Whitfield, 2002). The core oligosaccharide itself can be further split into an inner and outer core. The inner core is composed of up to three *L*-glycero-α-*D*-manno-heptose (Hep) residues, one of which is attached to a core 3-deoxy-*D*-manno-oct-2-ulosonic acid (Kdo) residue (Heinrichs *et al*., 1998) which may also have additional linked Kdo residues. The outer core is a non-repeating polysaccharide consisting of hexose residues attached to the heptosyl backbone (Raetz and Whitfield, 2002). hSP-D recognises bacterial LPS through association with heptose and the LPS inner core (Kuan *et al*., 1992; Wang *et al*., 2008; Clark *et al*., 2016), and this interaction varies with length, complexity and composition of the LPS.

A trimeric recombinant fragment of hSP-D (rfhSP-D) which has anti-inflammatory activity in murine models of inflammation (Strong et al., 2002; Clark et al., 2003; Knudsen et al., 2007) has provided key insights into bacterial recognition by hSP-D. The gram-negative bacteria *Haemophilus influenzae* and *Salmonella enterica* are of particular interest because while they can precipitate serious illness in both children and immune-compromised adults, they are also both recognised and neutralised by hSP-D (Murphy *et al*., 2009; Chattopadhyay *et al*., 2010). For example, *H. influenzae* serotype b strains are associated with invasive bacterial infections including meningitis and septicaemia (Moxon and Vaughn, 1981; Moxon and Maskell, 1992), whereas acapsular or non-typeable strains of *H. influenzae* can cause sinusitis, conjunctivitis, otitis media, and acute lower respiratory tract infections (Murphy *et al*., 2009). *S. enterica* is a pathogen that causes periodic outbreaks of gastroenteritis with some strains, such as *S. enterica* serovar (sv) Typhi, causing enteric fever (Coburn *et al*., 2007).

Crystallographic analyses have revealed how hSP-D interacts with, and recognises, a range of carbohydrate structures from relatively simple carbohydrates (Shrive *et al*., 2003; Crouch *et al*., 2006; Crouch *et al*., 2007; Wang *et al*., 2008; Shrive *et al*., 2009), to more complex structures, such as the purified core oligosaccharide from rough strains of the gram-negative bacteria *H. influenzae* and *S. enterica* (Clark *et al*., 2016; Littlejohn *et al*., 2018). In each case, recognition of carbohydrate by hSP-D is achieved through calcium-dependent binding, at the primary calcium site, of a mannose-type or stereochemically-equivalent equatorial hydroxyl pair such as O6⍰, O7⍰ of a core heptose or O3⍰, O4⍰ of a terminal glucose, with the binding pocket flanking residues Arg343 and Asp325 contributing to recognition and binding (Shrive *et al*., 2003; Crouch et al., 2009; Shrive *et al*., 2009; Clark *et al*., 2016; Littlejohn *et al*., 2018). The high-resolution structures published in Clark *et al*. (2016) and Littlejohn *et al*. (2018) clearly demonstrated that hSP-D specifically and preferentially targets the bacterial LPS inner core of *H. influenzae* Eagan and *S. enterica* sv Minnesota rough strains R5 and R7 (rough mutant chemotypes Rc and Rd1) via the innermost heptose residue. These ligand-bound structures also highlighted that hSP-D has the flexibility and versatility to recognise alternative LPS epitopes, for example a terminal glucose O3⍰, O4⍰ pair, when the preferred core heptose is unavailable (Littlejohn *et al*., 2018).

In order to separate the core oligosaccharide from lipid A for use in protein crystallisation studies, bacterial LPS is hydrolysed with acetic acid (Masoud, *et al*., 1997). While this efficiently separates the hydrophilic core oligosaccharide from the hydrophobic lipid A domain, which is not involved in hSP-D recognition (Kuan et al., 1992; Lim et al., 1994), acid hydrolysis often also cleaves not only functional groups such as phosphate attached to core oligosaccharide residues but also Kdo residues extending from the central core Kdo. The β-elimination of Kdo-linked phosphate leads to the reorganisation of Kdo into a 5-membered furanoid derivative, otherwise known as an anhydro-Kdo (Auzanneau *et al*., 1991; Clark *et al*., 2016). Accordingly, it has not been possible via this route to visualise the interaction of rfhSP-D with an intact Kdo residue, nor has it been possible to reliably establish the role in ligand recognition by rfhSP-D of functional groups such as phosphate which are attached to core oligosaccharide residues. Previous binding and modelling studies of the interaction of SP-D with synthetic LPS inner core structures (Reinhardt *et al*., 2016), using glycan array and surface plasmon resonance measurements, included non-phosphorylated representative *H. influenzae* and *S. enterica* core LPS components including the *H. influenzae* inner core HepIII-(1,2)-HepII-(1,3)-HepI and the HepI-(1,5)-KdoI common to both bacteria. These studies suggested that the Kdo does not influence binding but that the outer core residues might affect the preferred binding via HepI (Littlejohn *et al*., 2018).

The work presented here builds on our previous research by using reported synthetic di- and trisaccharides (Ekelöf and Oscarson, 1995; Ekelöf, 1996; Bernlind and Oscarson, 1997), which correspond to elements of the core oligosaccharides of many gram-negative bacteria including *H. influenzae* and *S. enterica*, to overcome the disadvantageous effects of acid hydrolysis on bacterial LPS purified from whole bacteria. As a result, this work has allowed us to explore structurally for the first time not only how rfhSP-D recognises the intact HepI-Kdo element of the LPS core (Reinhardt *et al*., 2016) but also how rfhSP-D interacts with variably phosphorylated inner core oligosaccharide fragments. Ultimately, this promotes a greater understanding of how proteins of the innate immune response, in this case hSP-D and the biologically and therapeutically active trimeric recombinant fragment rfhSP-D, can recognise and interact with clinically important human pathogens.

## RESULTS

Four synthetic oligosaccharides based on inner core motifs of *H. influenzae* Eagan and *S. enterica* sv Minnesota rough strains LPS were used for soaking into native crystals of rfhSP-D (see Figure 1). These oligosaccharides all contain the inner core heptose that has been shown to be preferentially bound to hSP-D and also include a spacer on the reducing terminal sugar, replacing as appropriate where the heptose-linked inner core Kdo or lipid A would be attached. Three of these oligosaccharides, the triheptoside structure (HepIII-(1,2)-HepII-(1,3)-HepI) (Bernlind and Oscarson, 1997) and the two phosphorylated disaccharides (HepII-(1,3)-HepI-4-PhosI and PhosII-4-HepII-(1,3)-HepI) (Ekelöf and Oscarson, 1995) include a p-trifluoracetamidophenylethyl spacer while the Hep-Kdo disaccharide (HepI-(1,5)-KdoI) (Ekelöf, 1996), common to both bacteria, includes a p-aminophenylethyl analogue (see Figure 1).

**Figure 1.**
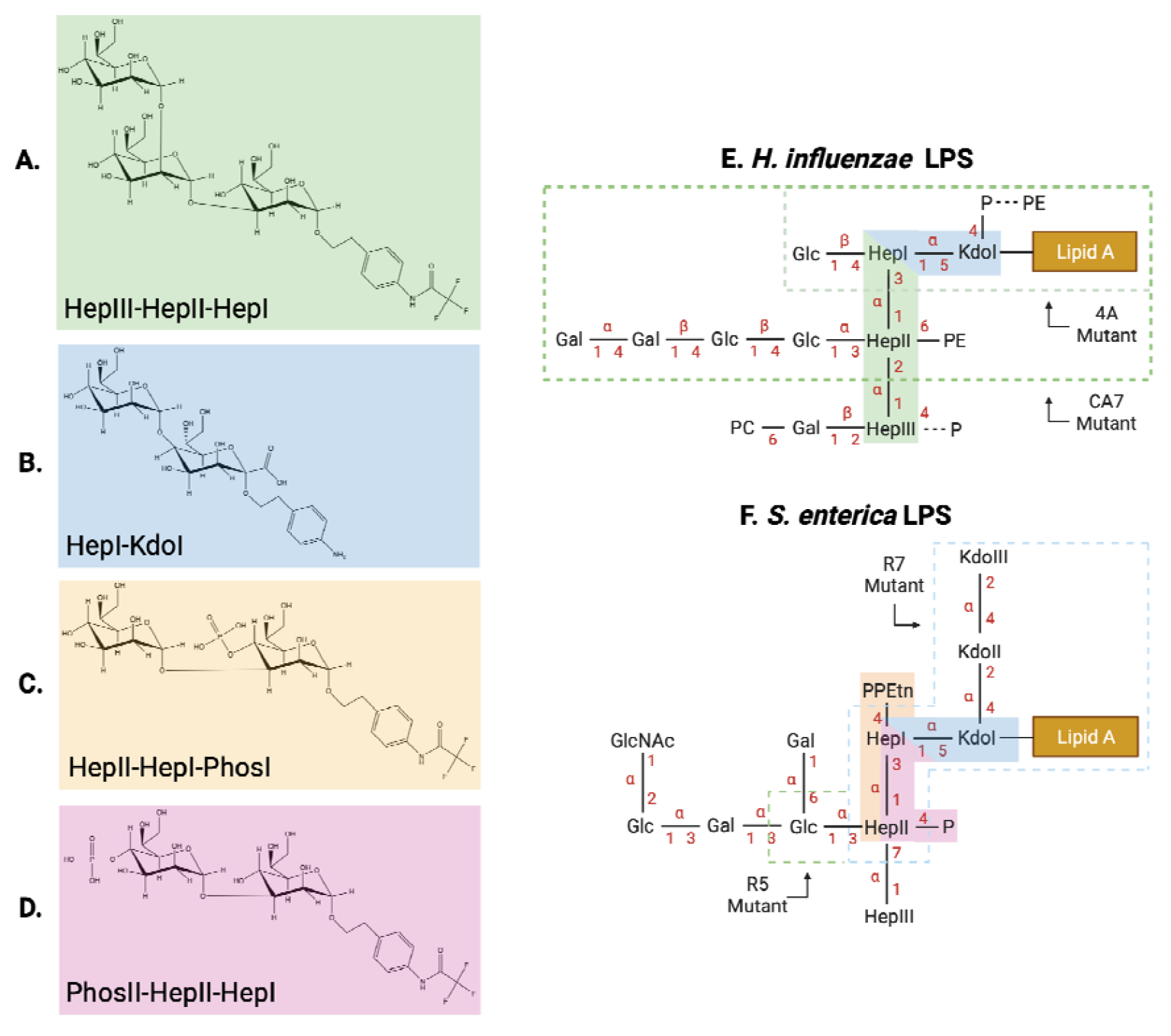
Structure of the synthetic oligosaccharides corresponding to inner core elements of LPS from *H. influenzae strain Eagan and S. enterica sv Minnesota*. **(A)** The HepIII-(1,2)-HepII-(1,3)-HepI trisaccharide (Bernlind and Oscarson, 1997) which maps on to the *H. influenzae* LPS inner core (**B**) The HepI-(1,5)-KdoI disaccharide (Ekelöf, 1996) which maps on to the inner core of both *H. influenzae* and *S. enterica* LPS (**C**) HepII-(1,3)-HepI-4-PhosI and (**D**) PhosII-4-HepII-(1,3)-HepI (Ekelöf and Oscarson, 1995) which both map on to the *S. enterica* LPS inner core. (**E**) The structure of LPS isolated from wild-type *H. influenzae* Eagan, adapted from Hood *et al*. (1996) with 4A and CA7 referring to the rfaF and orfH mutant strains, respectively. (**F**) The structure of LPS isolated from *S. enterica* rough mutant strains with the Rc(R5) and Rd1(R7) phenotypes indicated. LPS structures adapted from Mansfield and Forsythe (2001). Glc, glucose; GlcNAc, N-acetyl-glucosamine; Gal, galactose; Hep, L-D-Heptose; P, phosphate; PE/ PEtn, phosphoethanolamine; PC, phosphocholine; Kdo, 3-deoxy-D-manno-oct-2-ulosonic acid. Chemical depictions of synthetic oligosaccharides produced in ChemDraw. Figure generated using BioRender.

Crystal structures of rfhSP-D in complex with each of the synthetic oligosaccharides have been determined to high resolution, as follows: HepI-(1,5)-KdoI (1.75 Å), HepIII-(1,2)-HepII-(1,3)-HepI (1.63 Å), HepII-(1,3)-HepI-4-PhosI (1.92 Å), and PhosII-4-HepII-(1,3)-HepI (1.85 Å). The structures reveal ligand bound at the Ca1 binding pocket in two subunits (B and C) of the rfhSP-D trimer (Supplementary Figure S1) and no bound ligand in the third (subunit A) consistent with the tight crystal packing around the ligand-binding pocket, as previously reported (Clark *et al*., 2016). Protein and ligand-calcium distances for each of the structures are provided in Table 1 and Supplementary Table 1.

**Table 1.**
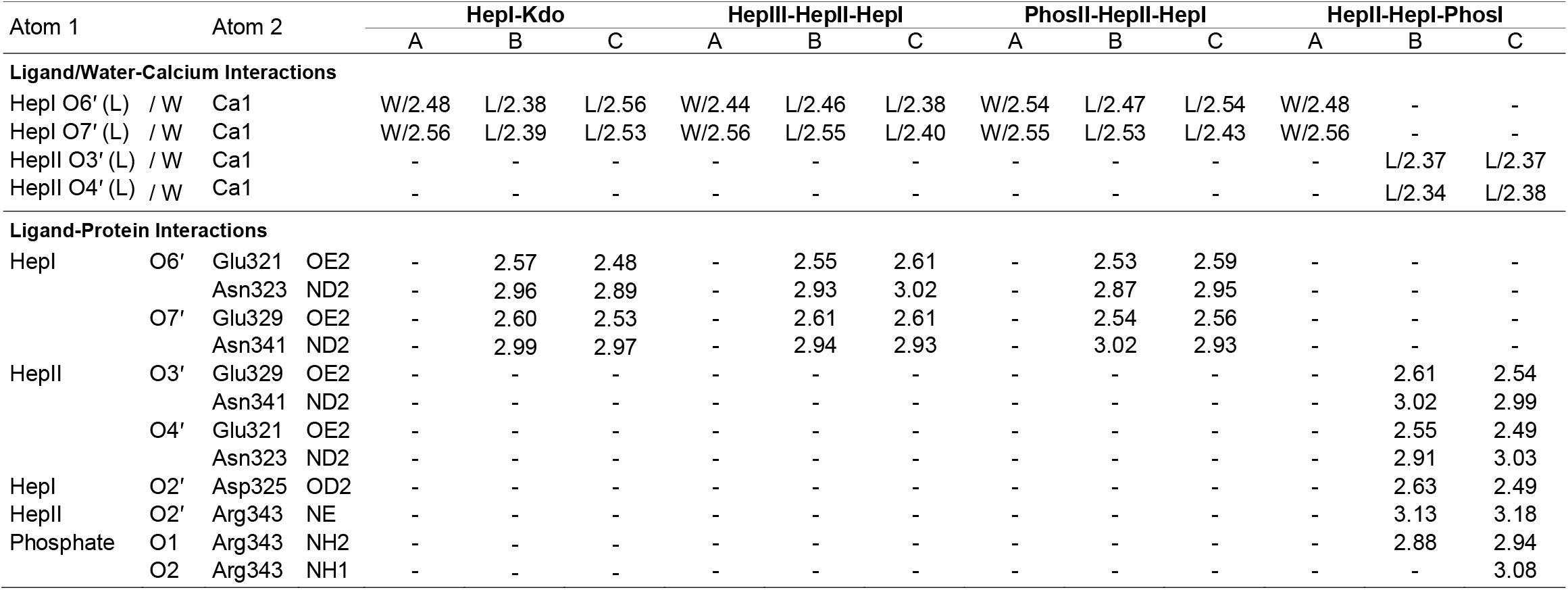
Ligand (L)/Water (W) – Calcium/Protein Bond Lengths (Å).

| Atom 1 |  | Atom 2 |  | HepI-Kdo |  |  | HepIII-HepII-HepI |  |  | PhosII-HepII-HepI |  |  | HepII-HepI-PhosI |  |  |
| --- | --- | --- | --- | --- | --- | --- | --- | --- | --- | --- | --- | --- | --- | --- | --- |
|  |  |  |  | A | B | C | A | B | C | A | B | C | A | B | C |
| Ligand/Water-Calcium Interactions |  |  |  |  |  |  |  |  |  |  |  |  |  |  |  |
| HepI | O6' (L) / W | Ca1 |  | W/2.48 | L/2.38 | L/2.56 | W/2.44 | L/2.46 | L/2.38 | W/2.54 | L/2.47 | L/2.54 | W/2.48 | - | - |
| HepI | O7' (L) / W | Ca1 |  | W/2.56 | L/2.39 | L/2.53 | W/2.56 | L/2.55 | L/2.40 | W/2.55 | L/2.53 | L/2.43 | W/2.56 | - | - |
| HepII | O3' (L) / W | Ca1 |  | - | - | - | - | - | - | - | - | - | - | L/2.37 | L/2.37 |
| HepII | O4' (L) / W | Ca1 |  | - | - | - | - | - | - | - | - | - | - | L/2.34 | L/2.38 |
| Ligand-Protein Interactions |  |  |  |  |  |  |  |  |  |  |  |  |  |  |  |
| HepI | O6' | Glu321 | OE2 | - | 2.57 | 2.48 | - | 2.55 | 2.61 | - | 2.53 | 2.59 | - | - | - |
|  |  | Asn323 | ND2 | - | 2.96 | 2.89 | - | 2.93 | 3.02 | - | 2.87 | 2.95 | - | - | - |
|  | O7' | Glu329 | OE2 | - | 2.60 | 2.53 | - | 2.61 | 2.61 | - | 2.54 | 2.56 | - | - | - |
|  |  | Asn341 | ND2 | - | 2.99 | 2.97 | - | 2.94 | 2.93 | - | 3.02 | 2.93 | - | - | - |
| HepII | O3' | Glu329 | OE2 | - | - | - | - | - | - | - | - | - | - | 2.61 | 2.54 |
|  |  | Asn341 | ND2 | - | - | - | - | - | - | - | - | - | - | 3.02 | 2.99 |
|  | O4' | Glu321 | OE2 | - | - | - | - | - | - | - | - | - | - | 2.55 | 2.49 |
|  |  | Asn323 | ND2 | - | - | - | - | - | - | - | - | - | - | 2.91 | 3.03 |
| HepI | O2' | Asp325 | OD2 | - | - | - | - | - | - | - | - | - | - | 2.63 | 2.49 |
| HepII | O2' | Arg343 | NE | - | - | - | - | - | - | - | - | - | - | 3.13 | 3.18 |
| Phosphate | O1 | Arg343 | NH2 | - | - | - | - | - | - | - | - | - | - | 2.88 | 2.94 |
|  | O2 | Arg343 | NH1 | - | - | - | - | - | - | - | - | - | - | - | 3.08 |

The extent to which each ligand can be modelled into the electron density varies between structures due to the nature and extent of contact and interaction with the protein. In the HepI-(1,5)-KdoI bound structure HepI is clearly defined in the electron density, but there is no evidence to support the fitting of the Kdo residue or the spacer (Figure 2A). For the HepIII-(1,2)-HepII-(1,3)-HepI bound structure (Figure 2B), clear electron density is present for HepI – Hep II, and for the HepI-linked spacer in subunit B, but not for HepIII. For the phosphorylated ligands corresponding to *S. enterica* core components, the complete HepII-(1,3)-HepI-4-PhosI bound ligand, except for the spacer, is fully defined in both subunits (Figure 2C) while the whole of the PhosII-4-HepII-(1,3)-HepI ligand can be fitted into the subunit B electron density (Figure 2D). In subunit C there is only electron density present for HepI.

**Figure 2.**
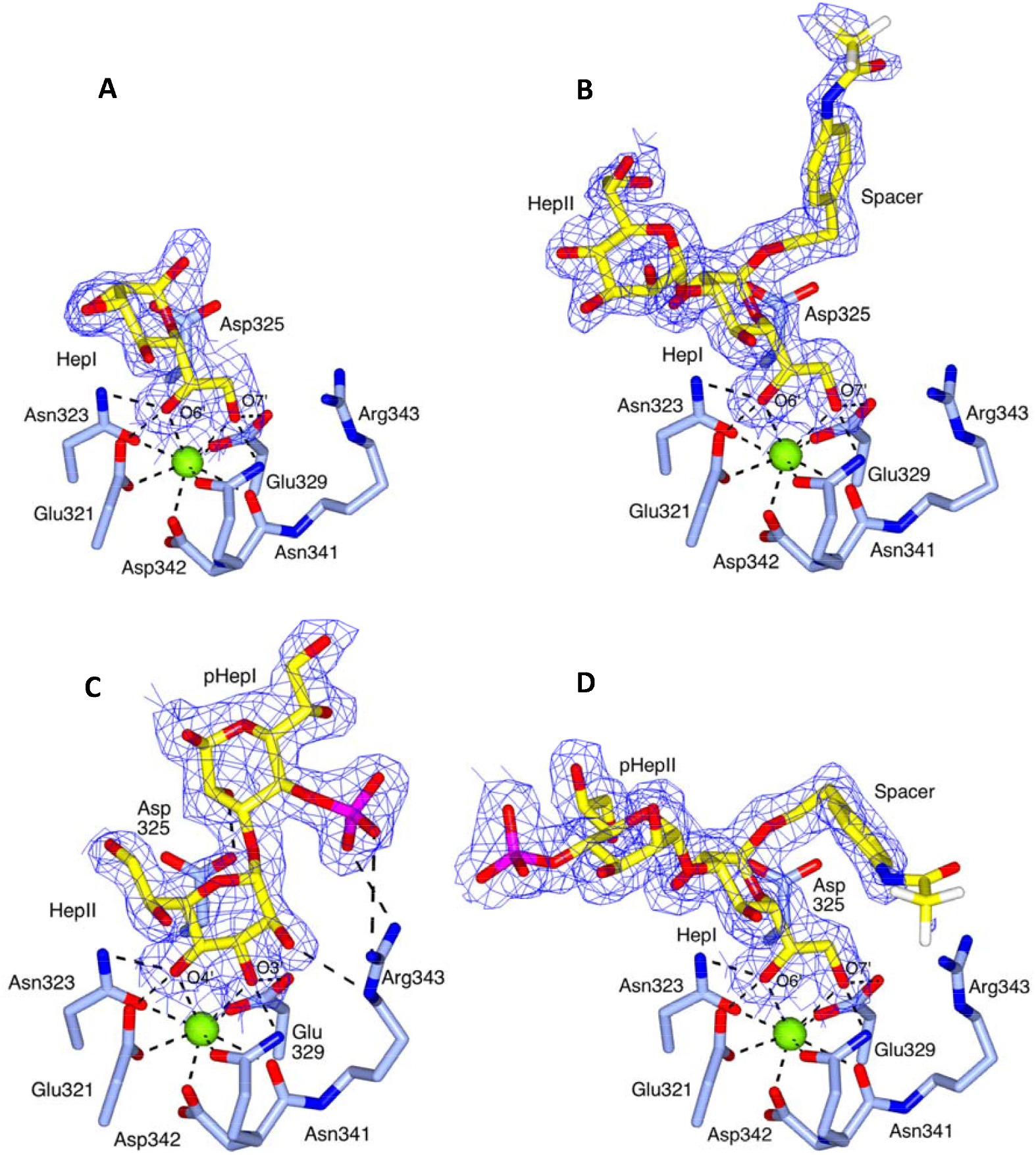
Coordination of the synthetic saccharides in the rfhSP-D subunit B Ca1 binding pocket. In each case the ligand (yellow) is coordinated to Ca1 (green sphere) with key interactions (dashed lines) and binding site residues (ice blue) indicated. The 2mFo-DFc electron density map (blue) is clipped to the bound ligand and contoured at 1σ. (**A**) HepI-(1,5)-KdoI. Only HepI, coordinated to Ca1 by the O6⍰ and O7⍰ hydroxyls, is visible in the electron density (**B**) HepIII-(1,2)-HepII-(1,3)-HepI with HepI, HepII and the spacer visible in the electron density and coordination to Ca1 via the HepI O6⍰ and O7⍰ hydroxyls (**C**) HepII-(1,3)-HepI-4-PhosI showing the alternative mode of recognition via the HepII O3⍰ and O4⍰ hydroxyls of HepII. The spacer linked to HepI is not visible in the electron density. (**D**) PhosII-4-HepII-(1,3)-HepI, coordinated to Ca1 by the HepI O6⍰ and O7⍰ hydroxyls. Figure created using CCP4mg.

Recognition of the HepI-(1,5)-KdoI, (HepIII-(1,2)-HepII-(1,3)-HepI, and PhosII-4-HepII-(1,3)-HepI ligands is achieved by Ca1 coordination of HepI O6⍰ and O7⍰ side chain hydroxyls (Table 1; Figure 2), while recognition of the HepII-(1,3)-HepI-4-PhosI ligand is achieved by Ca1 coordination of the HepII O3⍰ and O4⍰ ring hydroxyls (Table 1; Figure 2C). In all cases the ligands are also coordinated by the protein through these hydroxyl pairs (Shrive et al., 2003; Wang et al., 2008), by Glu321, Asn323, Glu329 and Asn341. For the HepII-(1,3)-HepI-4-PhosI ligand there are additional protein-ligand interactions (Table 1; Figure 2C), between Arg343 and both the HepII O2⍰ hydroxyl (NE, 3.13, 3.18 Å) and the HepI-linked phosphate (NH1, 3.08 Å; NH2, 2.88, 2.94 Å), and between Asp325 OD2 and HepI O2⍰ (2.49, 2.63 Å). In each structure, ligand coordination is supported by an extensive network of water-mediated contacts and some limited interactions with residues in symmetry-related molecules (Supplementary Table S2).

The ligands used in these experiments are each conjugated to a chemical spacer (replacing either the core Kdo or, for the HepI-(1,5)-KdoI ligand, where the lipid A GlcN would be attached to the Kdo). Electron density was only sufficiently well-defined to allow fitting of the spacer in subunit B of the HepIII-(1,2)-HepII-(1,3)-HepI (Figure 2B) and PhosII-4-HepII-(1,3)-HepI (Figure 2D) bound structures. The spacers do not interact directly with the coordinating protein but do form weak (3.3 – 3.5 Å) crystal packing and water-mediated interactions. Comparison of the spacer in the PhosII-4-HepII-(1,3)-HepI and the HepIII-(1,2)-HepII-(1,3)-HepI bound structures reveals a 90° rotation about the spacer C6 – C7 bond, suggesting that there are two alternate conformations (see Figure 3A).

**Figure 3.**
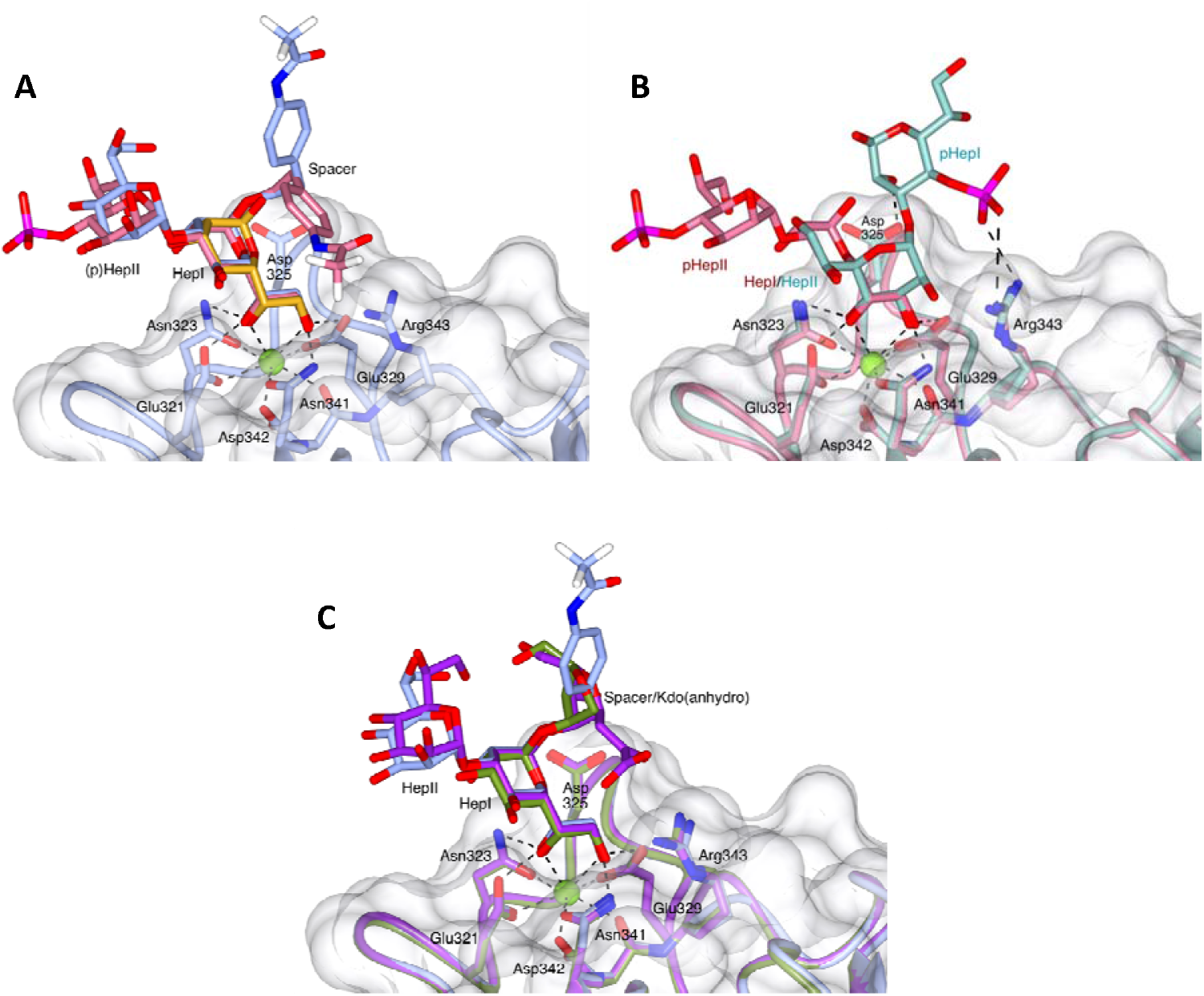
Overlays of the ligand bound structures and structures with bound anhydro Kdo. All overlays generated by a least-squares fit of subunit B main-chain atoms, with key interactions (dashed lines) and binding site residues, including Arg343 and Asp325, indicated. (**A**) HepI-(1,5)-KdoI (gold, only HepI visible in the electron density), HepIII-(1,2)-HepII-(1,3)-HepI-spacer (ice blue, HepIII not visible in the electron density) and PhosII-4-HepII-(1,3)-HepI-spacer (pale red). The two conformations of the HepI-linked spacer can be seen. The protein surface, main chain and side chains are of the HepIII-(1,2)-HepII-(1,3)-HepI bound structure. (**B**) The two phosphate substituted diheptosyl ligands showing the different binding modes and orientations of the two ligands. PhosII-4-HepII-(1,3)-HepI (pale red, HepI-linked spacer omitted for clarity) coordinated to Ca1 by the HepI O6⍰ and O7⍰ hydroxyls and HepII-(1,3)-HepI-4-PhosI (dark cyan, HepI-linked spacer not visible in the electron density) showing the alternative recognition mode via the HepII O3⍰ and O4⍰hydroxyls. The protein surface is of the PhosII-4-HepII-(1,3)-HepI structure; main chain and side chains are of both structures. (**C**) HepIII-(1,2)-HepII-(1,3)-HepI-spacer (ice blue, HepIII not visible in the electron density), *H. influenzae* Eagan 4A Hep-anhydro Kdo (olive green, PDB 4E52) and *S. enterica* sv Minnesota R7 HepII-HepI-anhydro Kdo (purple, PDB 5OXR). The protein surface is of the HepIII-(1,2)-HepII-(1,3)-HepI structure; main chain and side chains are of all three structures. Figure created using CCP4mg.

## DISCUSSION

The work presented here seeks to build on our current understanding of innate immune recognition of LPS by hSP-D (Clark *et al*., 2016; Littlejohn *et al*., 2018) through further structural studies of the interaction of hSP-D with the core LPS components of the two clinically important microbial species, *H. influenzae* and *S. enterica*. The LPS ligands used in previous structural studies of recognition by hSP-D of both *H. influenzae* (Clark *et al*., 2016) and *S. enterica* (Littlejohn *et al*., 2018) were obtained by acid hydrolysis which results in cleavage of functional groups attached to core oligosaccharide sugars by β-elimination and reorganisation of the core Kdo residue into an anhydro-furanoid derivative (Auzanneau *et al*., 1991; Clark *et al*., 2016). In the case of *S. enterica* (Littlejohn *et al*., 2018) this β-elimination also removed KdoII and KdoIII from the LPS (see Figure 1). While those studies revealed variable interactions of this anhydro-Kdo with the Ca1 binding pocket residues Asp325 and Arg343 and the ability of hSP-D to recognise alternative sugars when the preferred inner core is not accessible, reorganisation of the Kdo and the loss of acid labile groups such as phosphates (Phillips *et al*., 1992) precluded evaluation of hSP-D recognition of an unmodified Kdo and of the influence of key functional groups lost during hydrolysis. To address this, we used synthetic LPS components corresponding to the LPS inner core of *S. enterica, H. influenzae* and many other gram-negative bacteria (Figure 1). These ligands were soaked into native rfhSP-D crystals which gave high-resolution data (1.63 – 1.92 Å) revealing bound ligand (Figure 2).

In all the structures reported here, except for the HepII-(1,3)-HepI-4-PhosI LPS oligosaccharide component, ligand coordination is via the HepI O61/O71 hydroxyls to the calcium CRD and coordinating residues with no additional close, direct interactions with the protein surface. Water-mediated and crystal contacts do play a role in stabilising the ligand within the crystal, the latter preventing ligand access to the subunit A binding site. The orientation of the bound HepI is similar to that seen for heptose-bound hSP-D (Wang *et al*., 2008) and in other oligosaccharide structures (Clark *et al*., 2016; Littlejohn *et al*., 2018). For the HepI-bound ligands which include a second, α1-3 linked, heptose HepII, this second heptose is located in approximately the same direction with respect to HepI as that seen previously for hSP-D structures that contain HepII-(α1,3)-HepI bound via HepI (Wang *et al*., 2008; Clark *et al*., 2016; Littlejohn *et al*., 2018) (Figure 3A, 3C).

HepII-(1,3)-HepI linked carbohydrates are present in many core LPS structures, suggesting a key role for hSP-D recognition of HepI, with no direct involvement of HepII, in many gram-negative bacteria. This is supported by data for the HepIII-(1,2)-HepII-(1,3)-HepI trisaccharide, a conserved core oligosaccharide backbone of *H. influenzae* LPS (Figure 1) which suggests a clear preference for HepI recognition by hSP-D, also observed previously for *H. influenzae* strain Eagan 4A (Clark *et al*., 2016) and *S. enterica* Minnesota (Littlejohn *et al*., 2018). While extensions to the inner core heptosyl backbone can shield the core HepI residue from recognition by rfhSP-D (Clark *et al*., 2016), many *H. influenzae* strains, including non-typeable *H. influenzae* (NTHi) strains, such as NTHi strain 375, that are significant pathogens in children causing otitis media, sinusitis, and pneumonia (Murphy *et al*., 2009), produce LPS molecules lacking complex extensions to their core oligosaccharide backbone. It is the recognition of these particular bacterial LPS that the HepIII-(1,2)-HepII-(1,3)-HepI bound structure may help us understand further. Although the HepII-HepIII linkage in enterobacteria, such as *Salmonella*, is different (α1-7) to *Haemophilus*, the *S. enterica* Minnesota Rc (R5) and Rd1 (R7) mutants (Figure 1) have been shown to be bound via HepI in a similar manner (Littlejohn *et al*., 2018) suggesting that the nature of the HepII-HepIII linkage does not significantly influence the mode of HepI binding (see Figure 3C).

For the HepI-(1,5)-KdoI ligand, representing a core LPS component of many gram-negative bacteria (Figure 1B), the HepI residue is clearly present in the electron density, but there is no evidence for either the KdoI residue or the KdoI-linked spacer suggesting that there are either limited or no interactions between KdoI and hSP-D (Figure 2A). This supports the binding data obtained from glycan array analysis and surface plasmon resonance which shows little difference in the binding of wild-type hSP-D to Hep, Hep-(1-3)-Hep, and Hep-(1-5)-Kdo (Reinhardt *et al*., 2016). In the other ligand-bound structures here, the spacer is attached to the HepI where a Kdo would normally be found, but similar to KdoI of the HepI-(1,5)-KdoI ligand, the spacer makes no direct interactions with the protein surface even though different orientations of the spacer are seen for the two structures where the spacer is visible in the electron density (Figure 3C). Taken together, these data suggest that where HepI is (1,5) attached to a single Kdo residue, nominally KdoI, this Kdo sugar probably does not then have a significant role in ligand recognition. However, not all bacterial LPS structures contain a single Kdo residue, for example, *S. enterica* LPS contains up to three Kdo residues (Figure 1F). The question of whether the presence of these additional two Kdo residues, lost during acid hydrolysis of *S. enterica* LPS and hence not visible in the rfhSP-D LPS-bound structure (Littlejohn *et al*., 2018), might lead to interactions with the protein surface remains to be answered. The HepI-(1,5)-KdoI bound structure, where neither KdoI nor the spacer linked to KdoI where the lipid A would be attached are visible in the electron density, demonstrates how additional KdoI-linked Kdo residues, in for example Enterobacteria recognised by hSP-D (Kuan *et al*., 1992), and the lipid A, can be accommodated in the vicinity of the binding site.

The two phosphorylated ligands used in these experiments, which correspond to the HepI-HepII backbone in *S. enterica* LPS (Figure 1), are to the best of our knowledge the first structural definition of a human collectin molecule in complex with a phosphorylated oligosaccharide and differ only in the placement of the phosphate group, which is attached to O4⍰ of either HepI (HepII-(1,3)-HepI-4-PhosI) or HepII (PhosII-4-HepII-(1,3)-HepI). The HepII-(1,3)-HepI-4-PhosI LPS oligosaccharide component is the only ligand here that does not bind directly via HepI, recognition being achieved via the HepII O3⍰ and O4⍰ hydroxyls (Figure 2C). It is also the only ligand here that has additional interactions with the protein via the binding site flanking residues Arg343 and Asp325 (Table 1; Figure 2C). While the HepI O6⍰/O7⍰ hydroxyl pair on the HepII-(1,3)-HepI-4-PhosI ligand is technically available, the structure clearly shows that binding to SP-D in this manner would position the phosphate group unfavourably close to the protein surface, suggesting that when HepI is phosphorylated at O4⍰, binding via the HepI O6⍰/O7⍰ pair is no longer favourable, leading here to a reorientation of the ligand and binding via the HepII O3⍰/O4⍰ hydroxyls (Figure 3B). The core oligosaccharide of *S. enterica* LPS is only phosphorylated in longer, more complex structures, such as those found in *S. enterica* serovar Minnesota Ra mutant (Brade *et al*. 1985) (Figure 1) which, as illustrated by the structures presented here, is not recognised by hSP-D (Kuan *et al*., 1992) due to both phosphorylation of HepI and HepII at O4⍰ and the HepII O3’ pentasaccharide extension Glc-(Gal)Gal-Glc(GlcNAc) (Schnaitman and Klena, 1993). In the LPS from Rc (R5) and Rd1 (R7) mutants, which are bound by hSP-D (Kuan *et al*., 1992; Littlejohn *et al*., 2018), these phosphate substitutions are not present.

The structures presented here show that inner core HepI binding is the preferred mode of recognition with these small synthetic core oligosaccharides revealing no direct interactions between the protein and either an α1-3 linked HepII or KdoI. They also highlight the importance of Hep phosphorylation in both SP-D recognition and evasion, with phosphorylation of HepI at O4⍰ sterically preventing the preferred HepI O6⍰/O7⍰ binding, which is then overcome here by alternate recognition of the α1-3 linked HepII O3⍰/O4⍰ hydroxyl pair (Figure 3B). This alternative would not however be possible for inner core motifs which extend beyond HepII via an α1-3 link (see Figure 1), although other alternative modes of recognition, such as the terminal Glc recognition of *S. enterica* Minnesota R5 (Littlejohn *et al*., 2018), may then be adopted. Flexibility in LPS recognition appears to be a key attribute of the hSP-D protein. Previously, binding data from biochemical analyses and crystal structures have demonstrated that carbohydrate extensions to the core oligosaccharide can effectively shield inner core residues, for example HepI, from recognition by hSP-D (Clark *et al*., 2016). The structures presented here build on this concept by demonstrating that recognition is also significantly affected by phosphorylation of specific residues in the core LPS oligosaccharide. LPS phosphorylation is a consistent trend across gram-negative bacteria and likely provides an additional mechanism by which bacteria avoid detection by immune proteins, including hSP-D, that are encountered during their commensal and disease-causing lifestyles within the human host.

## METHODS

### Cloning, Expression, and Purification of rfhSP-D

The rfhSP-D protein was expressed in *Escherichia coli* using a bacterial expression system, as detailed in Madan *et al*., 2001 and Strong *et al*., 2002. Each chain contains 177 amino acids corresponding to residues Gly179 to Phe355, comprising a short (8 x Gly-Xaa-Yaa) collagen-like domain, an α-helical coiled-coil neck domain, followed by the CRD. The rfhSP-D was purified by a procedure involving denaturation and refolding of the inclusion bodies, followed by ManAc affinity and gel filtration chromatography. The rfhSP-D was passed through a polymyxin B column (Detoxi-Gel, Pierce) to remove endotoxin resulting in an endotoxin level of less than 10 pg/μg protein.

### Crystallisation and Data Collection

Native crystals of rfhSP-D were grown in sitting drops consisting of an equal volume of protein solution (1 μL) and precipitant solution, consisting of 16% PEG 4000 and 0.1 M Tris pH 8.0. Crystals were prepared for cryocooling using 2-methyl-2,4-pentanediol (MPD) in precipitant buffer. Ligand was introduced into the crystal by the addition of ligand to the cryobuffer. Successful additions of 2 – 3 μL aliquots of increasing concentrations (5-20%) of MPD cryobuffer were added to each well, followed by the addition of a further 2 μL aliquot of 20% MPD cryobuffer with a final exchange of 8 μL of the microbridge solution with 20% MPD cryobuffer. The concentration of each ligand in the respective cryobuffer was 20 mM, apart from the HepI-(1,5)-KdoI ligand, which was at a final concentration of 33.75 mM. Data were collected at Diamond Light Source stations I03 and I04, from a single crystal in each case, using a Pilatus3 6M detector. Integrated intensities were calculated using *Mosflm* (Battye *et al*., 2011) and data was processed using the *Aimless, Truncate, Uniquify*, and *Sortmtz* programmes as part of the CCP4 programme suite (Winn *et al*., 2011). Data collection and processing statistics are given in Table 2.

**Table 2.**
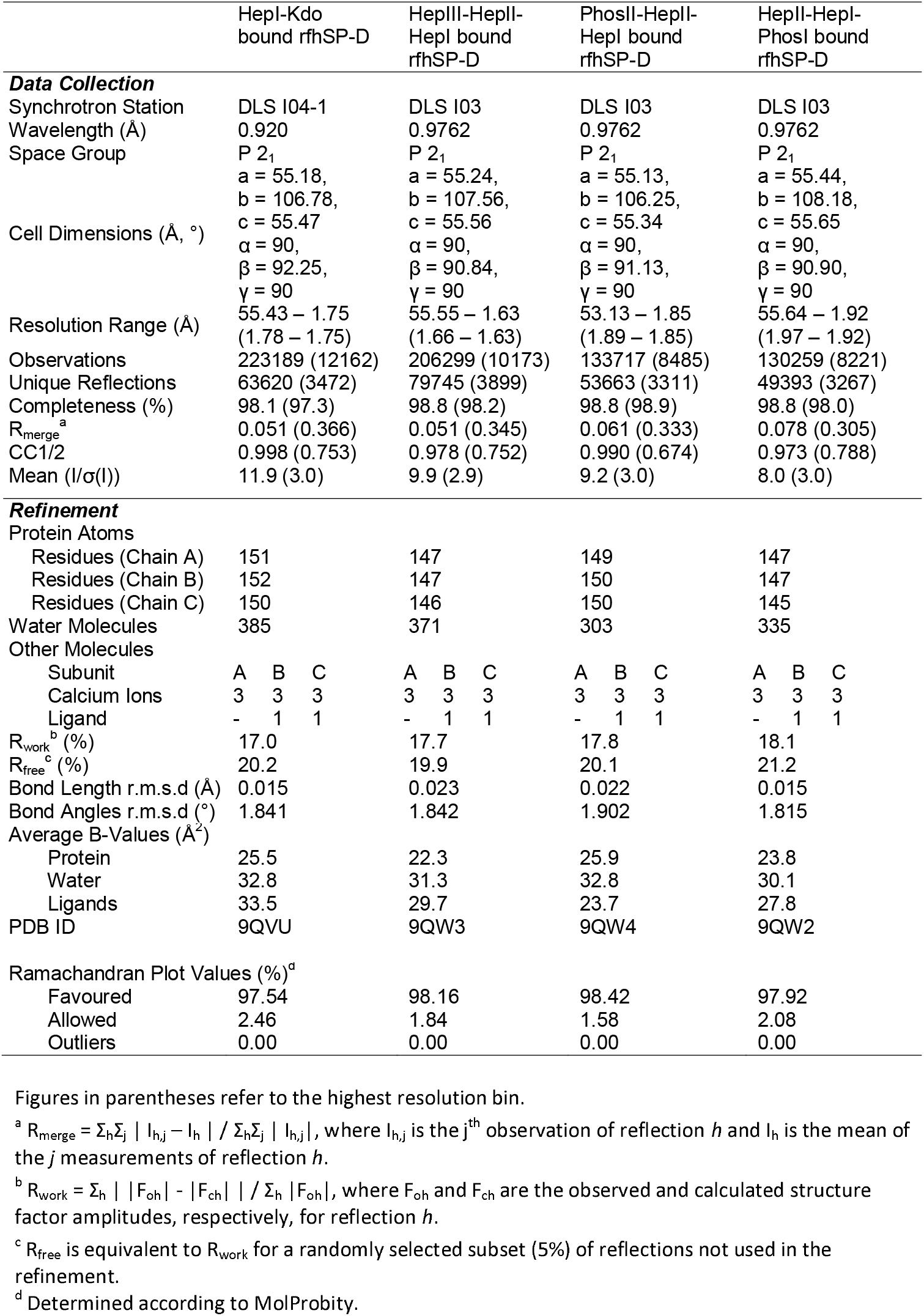
Crystallographic Data and Refinement Statistics.

### Structure Solution and Refinement

Isomorphism was sufficient to allow the atomic coordinates of the published 1.6 Å native rfhSP-D crystal structure (PDB: 1PW9, Shrive *et al*., 2003) to be used as a starting model for the rfhSP-D ligand-bound structures. Initial models were built in *Coot* (Emsley *et al*., 2010) after rigid body refinement with *Refmac5* (Murshudov *et al*., 2011). Subsequent model building was completed over multiple rounds of restrained refinement using *Refmac5* alternated with rounds of manual model building with *Coot*. Ligand coordinates were imported via *Coot* from the Protein Data Bank (PDB) with the exception of the two chemical spacers, which were generated manually using *AceDRG* (Long *et al*., 2017), which was also used to generate the dictionaries for the two chemical spacers. The quality of the final model was verified using the *MolProbity* server and *Privateer* as part of the CCP4i2 suite (Chen *et al*., 2010; Potterton *et al*., 2018) as well as the PDB validation software. Final refinement statistics are provided in Table 2. Molecular figures were generated using CCP4mg (McNicholas *et al*., 2011).

## ACKNOWLEDGEMENTS

We thank the beamline scientists at Diamond Light Source for their help and support. This work was supported by Science and Technology Facilities Council, and Diamond Light Source Awards MX14692 and MX19880 (TJG, AKS). S.O. was supported by Science Foundation Ireland (SFI) grants 08/IN.1/B2067 and 13/IA/1959.

J.M. and H.W.C would like to thank the Medical Research Council Catalyst for the Developmental Pathway Funding Scheme (DPFS) Award MR/P026907/1 and A.W. the University of Southampton MRC Doctoral Training Programme for funding his PhD.

## DATA AVAILABILITY

The coordinates and structure factors for the HepI-(1,5)-KdoI (9QVU), HepIII-(1,2)-HepII-(1,3)-HepI (9QW3), HepII-(1,3)-HepI-4-PhosI (9QW2) and PhosII-4-HepII-(1,3)-HepI (9QW4) ligand-bound structures have been deposited with the PDB alongside the coordinates and dictionaries for the two chemical spacers. The raw experimental diffraction image data are available via Keele University Data Repository (DOI: https://doi.org/10.21252/yzsx-0e16; DOI: https://doi.org/10.21252/nd1w-be36; DOI: https://doi.org/10.21252/m42p-9x07; DOI: https://doi.org/10.21252/bnzw-6034).

## CONFLICT OF INTEREST

The authors declare no conflicts of interest in regard to this manuscript.

## Notes

https://www.ebi.ac.uk/pdbe/entry/pdb/9qvu

https://www.ebi.ac.uk/pdbe/entry/pdb/9qw2

https://www.ebi.ac.uk/pdbe/entry/pdb/9qw3

https://www.ebi.ac.uk/pdbe/entry/pdb/9qw4

https://doi.org/10.21252/yzsx-0e16

https://doi.org/10.21252/nd1w-be36

https://doi.org/10.21252/m42p-9x07

https://doi.org/10.21252/bnzw-6034

### Competing Interest Statement

The authors have declared no competing interest.

