## Supplementary Tables and Figures for "Crystal structures of human SP-D complexed with synthetic oligosaccharides suggest a role for phosphorylated inner core LPS saccharides in host-pathogen interactions"

**Table S1.** Calcium-Protein/Water (W) Bond Lengths (Å).
**Table S2.** Interactions between ligands and residues in symmetry-related molecules (Å).
**Figure S1.** The rfhSP-D trimer with bound HepIII-HepII-HepI

**Table S1. Calcium-Protein/Water (W) Bond Lengths (Å).**

| Atom 1 | Atom 2 | | HepI-Kdo | | | HepIII-HepII-HepI | | | PhosII-HepII-HepI | | | HepII-HepI-PhosI | | |
|---|---|---|---|---|---|---|---|---|---|---|---|---|---|---|
| | | | A | B | C | A | B | C | A | B | C | A | B | C |
| Ca1 | Glu321 | OE1 | 2.58 | 2.55 | 2.48 | 2.57 | 2.52 | 2.63 | 2.50 | 2.50 | 2.73 | 2.56 | 2.48 | 2.68 |
| | Asn323 | OD1 | 2.44 | 2.43 | 2.40 | 2.36 | 2.35 | 2.37 | 2.45 | 2.35 | 2.47 | 2.41 | 2.53 | 2.51 |
| | Glu329 | OE1 | 2.50 | 2.40 | 2.36 | 2.43 | 2.42 | 2.42 | 2.38 | 2.31 | 2.38 | 2.35 | 2.41 | 2.39 |
| | Asn341 | OD1 | 2.39 | 2.44 | 2.40 | 2.36 | 2.39 | 2.39 | 2.40 | 2.34 | 2.43 | 2.34 | 2.37 | 2.36 |
| | Asp342 | O | 2.59 | 2.51 | 2.58 | 2.51 | 2.52 | 2.54 | 2.57 | 2.60 | 2.52 | 2.55 | 2.57 | 2.53 |
| | Asp342 | OD1 | 2.40 | 2.35 | 2.32 | 2.40 | 2.32 | 2.32 | 2.39 | 2.21 | 2.35 | 2.41 | 2.30 | 2.40 |
| Ca2 | Asp297 | OD1 | 2.66 | 2.58 | 2.64 | 2.55 | 2.62 | 2.59 | 2.62 | 2.67 | 2.76 | 2.62 | 2.66 | 2.63 |
| | Asp297 | OD2 | 2.42 | 2.42 | 2.58 | 2.44 | 2.44 | 2.43 | 2.38 | 2.59 | 2.37 | 2.40 | 2.48 | 2.40 |
| | Glu301 | OE1 | 2.50 | 2.47 | 2.47 | 2.40 | 2.47 | 2.45 | 2.50 | 2.42 | 2.55 | 2.52 | 2.53 | 2.57 |
| | Glu301 | OE2 | 2.56 | 2.52 | 2.55 | 2.47 | 2.48 | 2.48 | 2.49 | 2.55 | 2.54 | 2.47 | 2.53 | 2.51 |
| | Asn324 | OD1 | 2.52 | 2.58 | 2.59 | 2.55 | 2.65 | 2.56 | 2.60 | 2.44 | 2.54 | 2.51 | 2.48 | 2.54 |
| | Glu329 | O | 2.46 | 2.45 | 2.39 | 2.42 | 2.48 | 2.43 | 2.43 | 2.38 | 2.37 | 2.47 | 2.40 | 2.37 |
| | Asp330 | OD1 | 2.45 | 2.43 | 2.36 | 2.44 | 2.38 | 2.43 | 2.38 | 2.39 | 2.36 | 2.32 | 2.31 | 2.38 |
| | W | | 2.47 | 2.32 | 2.42 | 2.45 | 2.35 | 2.33 | 2.37 | 2.28 | 2.28 | 2.47 | 2.35 | 2.38 |
| Ca3 | Glu301 | OE1 | 2.28 | 2.40 | 2.36 | 2.42 | 2.37 | 2.37 | 2.32 | 2.35 | 2.46 | 2.28 | 2.46 | 2.31 |
| | Asp330 | OD1 | 2.50 | 2.57 | 2.58 | 2.54 | 2.52 | 2.57 | 2.60 | 2.48 | 2.60 | 2.56 | 2.54 | 2.60 |
| | Asp330 | OD2 | 2.46 | 2.51 | 2.48 | 2.45 | 2.50 | 2.48 | 2.50 | 2.53 | 2.47 | 2.42 | 2.40 | 2.52 |
| | W | | 2.30 | 2.18 | 2.24 | 2.20 | 2.26 | 2.33 | 2.14 | 2.27 | 2.25 | 2.00 | 2.29 | 2.23 |
| | W | | 2.36 | 2.37 | 2.38 | 2.29 | 2.38 | 2.35 | 2.27 | 2.35 | 2.31 | 2.27 | 2.27 | 2.26 |
| | W | | 2.38 | 2.37 | 2.45 | 2.36 | 2.40 | 2.35 | 2.32 | 2.40 | 2.37 | 2.38 | 2.32 | 2.35 |
| | W | | 2.58 | 2.38 | 2.45 | 2.37 | 2.40 | 2.38 | 2.41 | 2.43 | 2.46 | 2.51 | 2.51 | 2.39 |

**Table S2. Interactions between ligands and residues in symmetry-related molecules (Å).**

| Atom 1 | Atom 2 | HepI-Kdo | | | HepIII-HepII-HepI | | | PhosII-HepII-HepI | | | HepII-HepI-PhosI | | |
|---|---|---|---|---|---|---|---|---|---|---|---|---|---|
| | | A | B | C | A | B | C | A | B | C | A | B | C |
| HepI | O2′ Ser226/A OG | - | - | - | - | - | 2.85 | - | - | 2.91 | - | - | - |
| HepI | O2′ Ser226/C OG | - | 3.00 | - | - | 3.05 | - | - | - | - | - | - | - |
| HepII | O4′ Lys229/A NZ | - | - | - | - | | 2.54 | - | - | - | - | - | - |
| HepII | O4′ Lys229/C NZ | - | - | - | - | 2.61 | - | - | - | - | - | - | - |
| HepII | O7′ Ser226/C O | - | - | - | - | - | - | - | 2.76 | - | - | - | - |
| | O7′ Ser226/C OG | - | - | - | - | - | - | - | 3.09 | - | - | - | - |
| HepII | O6′ Tyr228/B OH | - | - | - | - | - | - | - | 3.17 | - | - | - | - |
| HepII/Phos | O1′ Tyr228/B OH | - | - | - | - | - | - | - | 2.59 | - | - | - | - |
| | O1′ Ser239/C OG | - | - | - | - | - | - | - | 2.62 | - | - | - | - |
| HepII/Phos | O2′ Lys229/C NZ | - | - | - | - | - | - | - | 2.91 | - | - | - | - |

**Figure S1. The rfhSP-D trimer with bound HepIII-HepII-HepI**.

The rfhSP-D trimer from the HepIII-(1,2)-HepII-(1,3)-HepI ligand-bound crystal structure is shown. Each protomer also binds three calcium ions, which are represented as green spheres. In subunits B (Gold) and C (Orange), where the HepIII-(1,2)-HepII-(1,3)-HepI ligand is bound, the ligand is shown at the Ca1 site in yellow. Note that in subunit B the HepI-linked spacer is visible in the map and was therefore fitted in the structure, in subunit C the spacer is missing. HepIII is not visible in the electron density in either subunit. Image created using CCP4mg.

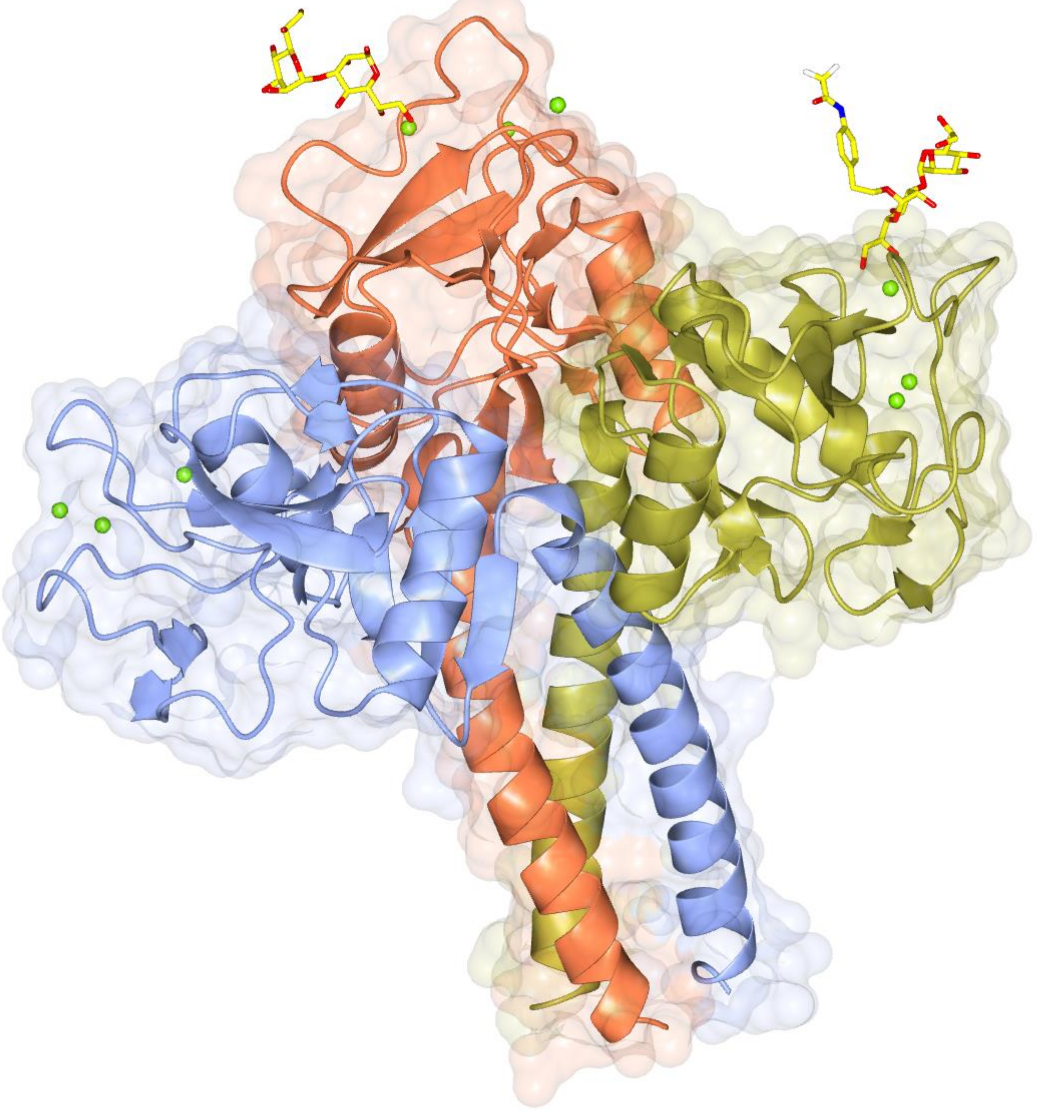
